# Small RNA sequencing of human sural nerves identifies widespread microRNA dysregulation and Schwann cell-localized miR-21-5p in diabetic peripheral neuropathy

**DOI:** 10.64898/2026.01.15.699768

**Authors:** Sneha A. Gummadi, Ryan R. Ju, Victoria Pastor, Eric C. Meyers, Shai M. Rozen, Diana Tavares-Ferreira

## Abstract

Diabetic peripheral neuropathy (DPN) is a common complication of diabetes with no disease modifying treatments. Despite the prevalence, the molecular mechanisms of DPN are not fully characterized. Among the various molecular regulators, microRNAs (miRNAs) control protein synthesis and are essential for normal development and homeostasis, with dysregulation implicated in cancer and neurodegenerative diseases. In this study, we performed small RNA-sequencing to profile the miRNA landscape of human sural nerves from individuals with and without DPN. Our analysis revealed that nearly 10% of all miRNAs detected are dysregulated and among those 74% are significantly downregulated in DPN. Target gene enrichment analysis of the differentially expressed miRNAs yielded pathways significantly associated with nerve regeneration, metabolic dysfunction, and immune cell activity. In particular, miR-21-5p is significantly upregulated in DPN, showed a positive association with axonal loss severity, and localizes to Schwann cells, consistent with its broader role as an injury- and inflammation-responsive miRNA that shifts from early pro-regenerative functions to maladaptive, inflammation-amplifying effects that impair Schwann cell mediated nerve repair. These results suggest that miRNAs may contribute to peripheral nerve degeneration by promoting inflammation, apoptosis, oxidative stress, and impaired nerve regeneration, while also opening potential avenues for biomarker discovery and therapeutic intervention.

**Article highlights:**

- We undertook this study to address the limited understanding of molecular changes contributing to diabetic peripheral neuropathy (DPN) in humans.
- We sought to profile microRNAs (miRNAs), key post-transcriptional regulators of gene expression, in human sural nerves and developed a dedicated computational pipeline for robust miRNA quantification, differential expression, and target enrichment analysis.
- Our analyses revealed widespread miRNA dysregulation in DPN, with most altered miRNAs downregulated and miR-21-5p significantly upregulated in DPN, highly correlated with axonal loss severity and localized to Schwann cells.
- These findings suggest that miRNA imbalance, including elevated Schwann cell miR-21-5p, may contribute to nerve dysfunction in DPN and provide new opportunities for biomarker development and therapeutic targeting.

## Introduction

Diabetes is a significant public health problem affecting an estimated 828 million adults worldwide ^1^. A common and debilitating complication is diabetic peripheral neuropathy (DPN) with nearly 50% to 66% of diabetic patients eventually developing it in their lifetime ^2^. Clinically, DPN typically presents with pain and/or diminished sensation ^3^. Patients may also develop wounds that fail to heal, increasing the risk for limb amputation and sepsis in severe cases ^3^.

Despite the prevalence and risks, the exact molecular mechanisms of DPN are not well understood. Type 2 diabetes is characterized by widespread metabolic dysfunction including hyperglycemia, dyslipidemia, insulin resistance, and mitochondrial dysregulation, all of which contribute to oxidative stress, inflammation, and progressive cellular damage ^4^. Peripheral nerves and their supporting Schwann cells, which play a vital role in maintaining the integrity and function of neurons, are highly susceptible to such damage due to their high metabolic demands ^5^. Given this susceptibility, mechanisms which regulate cellular homeostasis are of particular interest.

microRNAs (miRNAs) are small, non-coding RNAs, which are typically 18-25 nucleotides long that regulate gene expression ^6^. Canonically, miRNAs are transcribed as pri-miRNAs in the nucleus, trimmed by Drosha into pre-miRNAs that are exported to the cytoplasm, and then cleaved by Dicer to generate the mature miRNA ^6^. The mature miRNAs then form a complex with the RNA binding protein Argonaute2 (Ago2) to form an RNA-induced silencing complex (RISC), which bind to the 3’ untranslated region of a target messenger RNA (mRNA), either preventing mRNA translation or promoting mRNA degradation ^6^. The net effect of this process is decreasing the activity of specific genes and their subsequent biological process, allowing microRNAs to fine-tune molecular and cellular functions.

Previous studies have shown that reactive oxidative species (ROS), which are elevated in diabetes, cause transcriptional changes in miRNA expression ^7^. In addition, impaired miRNA maturation has been linked to worsened neuronal damage in DPN and other peripheral neuropathies ^8,9^. Furthermore, Schwann cell derived exosomes, which are enriched in miRNAs, have also been proposed as a potential therapy for DPN after reducing neuronal injury in rodent models ^10^. Beyond their regulatory roles, miRNAs such as hsa-miR-146 and hsa-miR-199a-5p have been investigated as candidate biomarkers of DPN ^11,12^.

Although dysregulation of miRNAs has been implicated in the mechanisms underlying DPN, a comprehensive characterization of their expression profiles in human peripheral nerves was lacking. In this study, we address this gap by profiling miRNA expression in human sural nerves from individuals with and without DPN and by examining how these miRNAs may regulate pathways involved in nerve regeneration, fibrosis, inflammation and immune activity.

## Methods

### Patient Consent

All protocols were reviewed and approved by the UT Dallas Institutional Review Board.

### Sural Nerve Tissue Collection and Processing

DPN sural nerve samples were recovered from patients (Table 1) undergoing lower limb amputation surgery for non-reconstructable soft tissue or bone, osteomyelitis, and/or critical limb ischemia as described previously ^13^. The tissue was then placed in sterile specimen cup, frozen with liquid nitrogen, and stored in −80C freezer. Control sural nerve samples were obtained from patients (Table 1) undergoing cross facial nerve grafting in the context of facial paralysis surgery. The sural nerve was harvested from a 3 cm vertical incision midway between the lateral malleolus and Achilles tendon. The nerve was then removed after cutting the proximal and distal ends of the needed length. The tissue was frozen immediately in pulverized dry ice, transferred to a sterile vial, and then stored in a −80C freezer.

**Table 1.** Patient Metadata.

| **Patient** | **Age** | **Sex** | **Diabetes** | **DPN** | **Pain vs. Numbness** | **History of Diabetic Foot Ulcers?** |
| --- | --- | --- | --- | --- | --- | --- |
| Control1 | 36 | Male | No | No | - | - |
| Control2 | 73 | Male | No | No | - | - |
| Control3 | 52 | Female | No | No | - | - |
| Control4 | 40 | Male | No | No | - | - |
| Control5 | 59 | Female | No | No | - | - |
| Control6 | 60 | Male | No | No | - | - |
| DPN1 | 31 | Male | Type 2 | Yes | Throbbing Pain | Yes |
| DPN2 | 79 | Male | Type 2 | Yes | Sharp Pain | Yes |
| DPN3 | 53 | Female | Type 2 | Yes | Numbness | Yes |
| DPN4 | 51 | Male | Type 2 | Yes | Numbness | Unknown |
| DPN5 | 59 | Female | Type 2 | Yes | Burning, Sharp Pain | Yes |
| DPN6 | 60 | Male | Type 2 | Yes | Constant Dull Pain, Sharp Pain | Yes |

### Small RNA Extraction, Library Preparation, and Sequencing

Purification of miRNA and total RNA was done following the protocol for Qiagen miRNAeasy Micro Kit (cat# 217084) for all 12 samples. Library preparation was completed using Nextflex Small RNA Library v4 per protocol and sequencing was done on NovaSeq X (Psomagen).

### Preprocessing and Alignment of Reads

Reads were trimmed using the cutadapt (v4.6) program using the following commands: -Z -a TGGAATTCTCGGGTGCCAAGG –minimum-length 16 --max-n 0 --length 35 and run again to perform quality trimming with the following commands: -Z --quality-cutoff 20 --minimum-length 16^14^. Next UniVec contaminants and ribosomal RNA (rRNA) were removed by exceRpt ^15^. Reads which passed UniVec and rRNA filtering completed the rest of the exceRpt protocol in the default alignment/mapping order for small RNAs: miRNA, tRNA, piwiRNA, gencode, and circRNA. The genome aligned bam file from exceRpt was run through HTSeq-count (v2.0.3) using the miRBase (v22.1) miRNA gff file to obtain the raw mature miRNA counts, which excludes ambiguous and non-unique reads for a given miRNA ^16,17^. The raw mature miRNA counts were then used for differential expression analysis.

### Differential Expression of miRNAs

Differential expression analysis was done using the DESeq2 package (v1.46.0) ^18^. Raw counts were provided as input, comparing miRNA expression of samples with and without DPN. Log fold shrinkage was performed using the apeglm function to reduce noise while maintaining large fold changes ^19^. Significant differential expression was defined by having an adjusted p-value < 0.05 and an absolute log2 fold change (LFC) > 0.58 (fold change > 1.5).

### Functional Analysis of Differentially Expressed miRNAs

Differentially expressed miRNAs were split into two groups: upregulated and downregulated miRNAs. Targets genes for each miRNA from miRTarBase 2025 and TargetScan 8.0 were extracted ^20–22^. Experimentally validated miRNA targets with strong evidence from miRTarBase were included. TargetScan predicted miRNA targets with a minimum weighted context++ percentile greater than 90 or maximum relative predicted K_d_ value of −3.0 were included. Only genes detected in bulk RNA-seq of sural nerve samples were included in the final miRNA target genes list ^13^. At least 5 miRNAs were required to target a respective gene for it to be considered for enrichment analysis. Enrichment analysis was conducted through clusterProfiler v4.12.6 using R v4.4.3 ^23,24^. Gene Ontology (GO), Reactome, WikiPathways, and Kyoto Encyclopedia of Genes and Genomes (KEGG) databases were used for analysis. An adjusted p-value < 0.05 using the Benjamini-Hochberg correction was considered significant for an enrichment term. Enrichment terms were filtered into the following categories: interleukins, MAPK, TGF-β/SMAD, PI3K/AKT, immune cell activity, AGE/RAGE, TNF-α, Neurotrophic factors, nerve regeneration, cell death, extracellular matrix/fibrosis, hyperglycemia/insulin/dyslipidemia, and miscellaneous. Inclusionary key terms were identified for each category (except miscellaneous), and additional exclusionary key terms for enrichment terms associated with organ systems unrelated to sural nerve tissue were identified (see **Suppl. File 1**). Any terms which did not meet the criteria for a group were added to the miscellaneous category.

### miRNA and axonal loss severity correlation analysis

miRNA expression was quantified from a separate subset of small RNA sequencing data and normalized to counts per million reads (CPM). CPM values were log2-transformed prior to analysis. Axonal density / loss severity was encoded as an ordinal variable (0 = normal, 1 = moderate, 2 = severe) as described previously ^13^. Associations between miRNA expression levels and axonal loss severity were assessed using Spearman rank correlation at the sample level. For visualization, individual sample values are shown with jitter to reduce overlap, and dashed lines indicate linear trend.

### MiRNA-mRNA integrative analysis

miRNA-mRNA integrative analysis was performed using the MIRit package ^25^ to identify putative regulatory interactions between differentially expressed miR-21-5p and its predicted mRNA targets, using a recently published mRNA dataset ^13^. Predicted targets were filtered using experimentally validated and high/very high confidence prediction databases, and miR-21-5p interactions were restricted to inversely correlated miRNA–mRNA pairs. Resulting target sets were used for downstream pathway and functional enrichment analyses.

### RNAscope *in situ* hybridization and immunohistochemistry (IHC)

RNAscope Plus smRNA-RNA HD (ACD) was performed according to manufacturer protocols with an optimized 10-second protease digestion. RNA integrity was verified using ACD positive and negative control probes, and probe and donor details are provided in Supplementary Tables S1. For dual RNAscope and immunohistochemistry, sections were blocked and incubated with neuronal and Schwann cell markers (Chicken Polyclonal Antibody to Peripherin, dilution 1:500, Encor Biotechnology, catalog number CPCA-Peri; Mouse Monoclonal Antibody to SOX10, dilution 1:100, Abcam, catalog number ab216020), followed by fluorescent secondary antibodies.

### Image acquisition

Sural nerve sections were imaged on an Olympus FV4000 confocal microscope at 60-100X magnification using standardized acquisition settings, with multiple artifact-free fields collected per section.

### Code and Data Availability

Computational methods are bundled into a pipeline, which is loosely based on the nf-core framework ^26^ and is publicly available on GitHub at utdal/miRNA-Analysis (https://github.com/utdal/miRNA-Analysis). All data is available in Suppl. Files. Raw and processed data will also be made publicly available.

## Results

### Small RNA landscape in human sural nerves

The study design and workflow are summarized in **Figure 1A** and the subject metadata are described in **Table 1 and Suppl. Table 1**. To investigate miRNA expression changes associated with DPN, we collected sural nerve samples from twelve subjects and extracted miRNA for small RNA-sequencing and data analysis. Six samples were obtained from individuals with type II diabetes and clinically diagnosed DPN undergoing lower limb amputation (4 males and 2 females; mean age of 55.5 ± 14.2 years), and six age and sex matched control samples were obtained from individuals undergoing cross facial nerve graft procedures (4 males and 2 females; mean age of 53.3 ± 12.5 years). We implemented an automated workflow for quality control, alignment, quantification, and downstream analysis of miRNA profiles. Across all samples, the vast majority of reads mapped to miRNAs, with smaller fractions aligning to tRNAs, piRNAs, circRNAs, rRNAs, and other annotated small RNAs (**Fig. 1B**). Out of 2,652 known miRNAs ^27^, 1,821 miRNAs were detected in at least one sample (count > 0).

**Figure 1.**
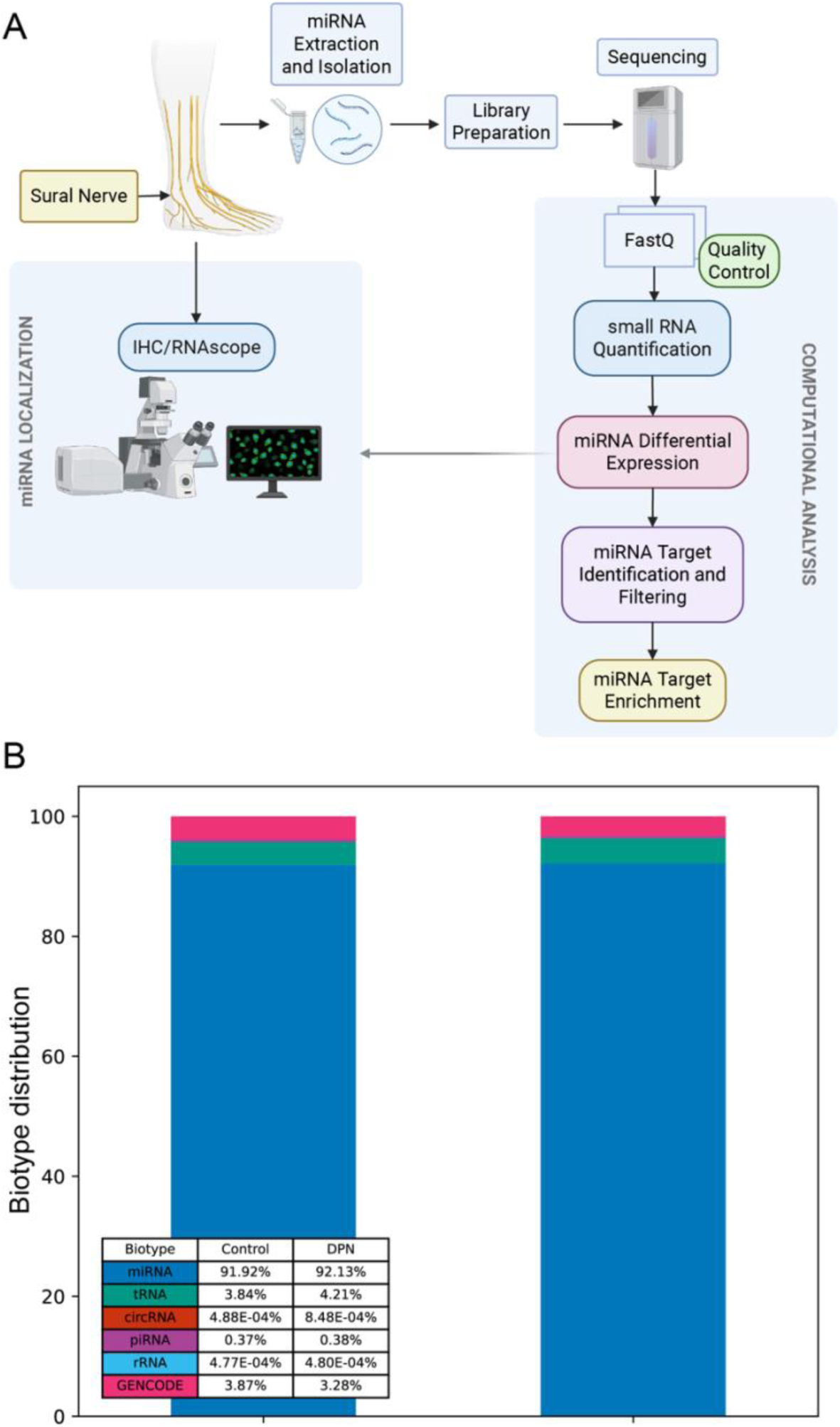
Overview. A) Overview of miRNA analysis and cell localization workflow. B) Distribution of different RNA biotypes across all samples using exceRpt.

The most abundant miRNAs across all samples were hsa-miR-23a-3p, hsa-miR-23b-3p, hsa-miR-143-3p, hsa-miR-126-3p, hsa-miR-99a-5p, hsa-miR-451a, hsa-miR-145-5p, hsa-miR-223-3p, hsa-let-7b-5p, and hsa-miR-21-5p (**Table 2**). Target enrichment analysis of the top ten miRNAs in DPN samples identified several genes that were recurrently regulated. Insulin-like Growth Factor 1 Receptor (*IGF1R*), N-alpha-acetyltransferase 50 (*NAA50*), Leucine Rich Repeat Containing 8 VRAC subunit B (*LRRC8BI*), Transforming Growth Factor-β Receptor 2 (*TGFBR2*), Homeobox A1 (*HOXA1*), and BCL2 apoptosis regulator (*BCL2*) were each predicted targets of at least five of the ten most highly expressed miRNAs, highlighting shared regulatory pressure on pathways related to growth signaling, cellular stress responses, and apoptosis.

**Table 2.** Top Expressed miRNAs in Control and DPN Sural Nerves.

| Control |  | DPN |  |
| --- | --- | --- | --- |
| miRNA | Mean | miRNA | Mean |
| hsa-miR-451a | 1811740.55 | hsa-miR-23a-3p | 1945302.69 |
| hsa-miR-23a-3p | 1692361.34 | hsa-miR-143-3p | 1745791.31 |
| hsa-miR-23b-3p | 1674226.83 | hsa-miR-23b-3p | 1716871.48 |
| hsa-miR-143-3p | 1427904.59 | hsa-miR-126-3p | 1295104.18 |
| hsa-miR-223-3p | 1301811.85 | hsa-miR-21-5p | 1007248.21 |
| hsa-miR-99a-5p | 961356.38 | hsa-miR-99a-5p | 1001781.18 |
| hsa-miR-126-3p | 843657.84 | hsa-miR-145-5p | 1001484.81 |
| hsa-miR-145-5p | 817329.00 | hsa-let-7b-5p | 761366.79 |
| hsa-let-7b-5p | 553180.61 | hsa-miR-27a-3p | 669514.13 |
| hsa-miR-27b-3p | 539095.45 | hsa-miR-100-5p | 631695.02 |

*Note.* Mean was calculated using DESeq2 normalized read counts.

### Widespread dysregulation of miRNA expression in human sural nerves

Next, we conducted differential miRNA expression analysis between DPN and control sural nerves. Principal component analysis showed separation between control and DPN samples, consistent with distinct miRNA expression profiles (**Fig. 2A**). Out of the 1,821 detected miRNAs, a total of 175 miRNAs (9.6%) were found to be differentially expressed between individuals with and without DPN at an adjusted p-value less than 0.05 and fold change (FC) higher than 1.5 (**Fig. 2B, Suppl. File 2**). Of those, 46 were upregulated and 129 were downregulated in DPN (**Fig. 2C**). Additionally, of the top 10 expressed miRNAs across all samples, hsa-miR-126-3p and hsa-miR-21-5p were found to be upregulated while hsa-miR-451a and hsa-miR-223-3p were found to be downregulated (**Fig. 2B**).

**Figure 2.**
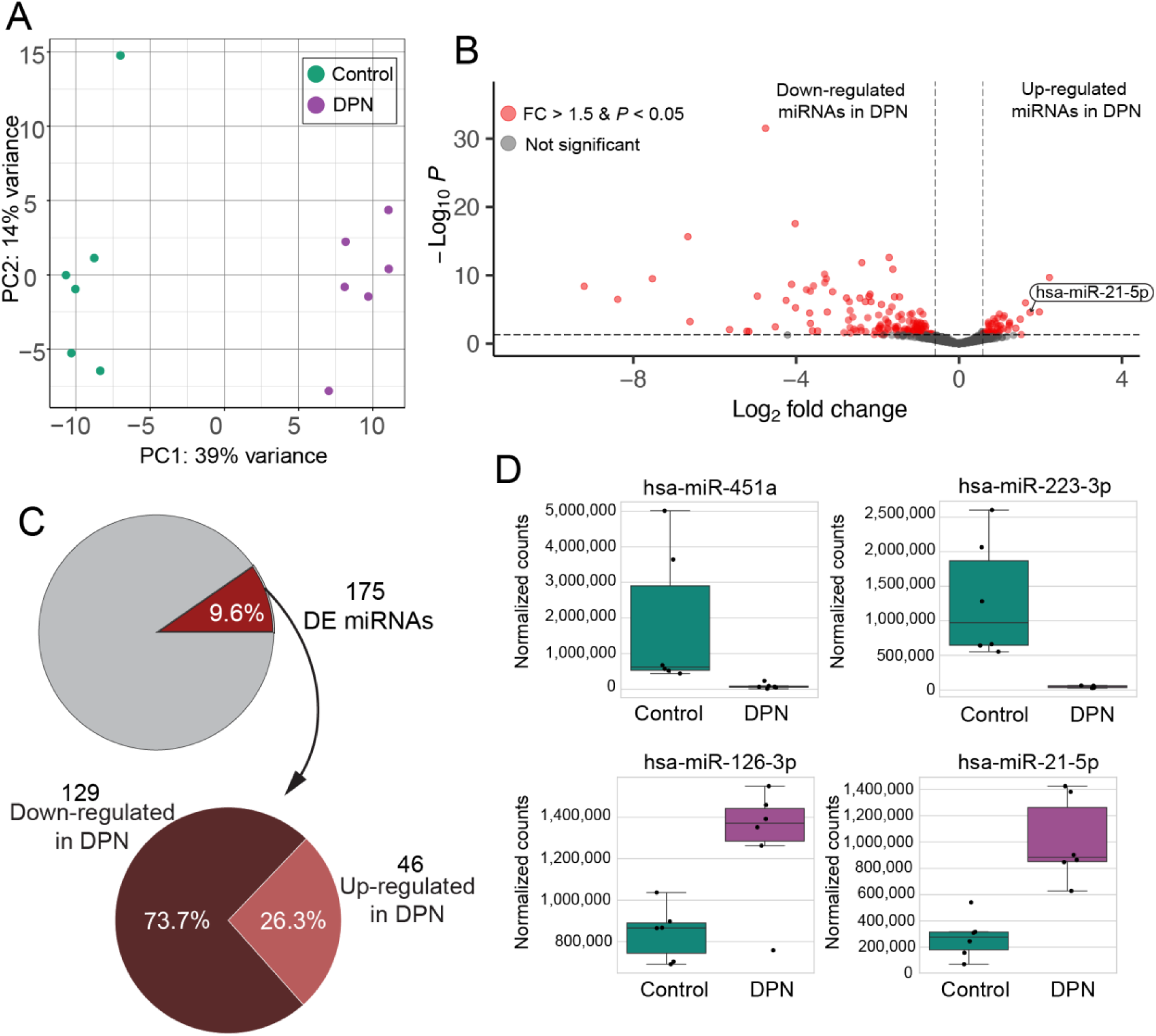
Small RNA-sequencing of human sural nerves reveals miRNA expression changes in DPN. A) Principal component analysis shows separation between control and DPN samples. B) Volcano plot showing differentially expressed genes in red (fold change > 1.5, adjusted P value < 0.05). C) Of the 1,821 detected miRNAs, nearly 10% were dysregulated. Among these, 129 (73.7%) were downregulated and 46 (26.3%) were upregulated in DPN relative to controls. D) Expression of the top expressed and differentially expressed miRNAs in DPN, hsa-miR-451a, hsa-miR-223-3p, hsa-miR-126-3p, and hsa-miR-21-5p in control and DPN samples. Counts are normalized by DESeq2.

Out of all the downregulated miRNAs in DPN, 35 showed complete loss of expression. Representative miRNAs include hsa-miR-7850-5p, hsa-miR-11401, hsa-miR-2115-5p, hsa-miR-2115-3p, and hsa-miR-6787-3p, all significantly reduced to near-undetectable levels in DPN samples, with a LFC < −6 (**Table 3**).

**Table 3.** miRNAs Downregulated to Nearly No Detection in DPN with LFC < −6.

| miRNA | Control Mean Count (±SE) | DPN Mean Count (±SE) | LFC |
| --- | --- | --- | --- |
| hsa-miR-7850-5p | 23.28 (±10.21) | 0 (±0) | -9.20 |
| hsa-miR-11401 | 14.79 (±5.16) | 0 (±0) | -8.38 |
| hsa-miR-2115-5p | 33.53 (±11.45) | 0.17 (±0.17) | -7.53 |
| hsa-miR-2115-3p | 94.42 (±27.72) | 0.62 (±0.40) | -6.66 |
| hsa-miR-6787-3p | 6.05 (±2.30) | 0 (±0) | -6.61 |

*Note.* Mean is calculated using DESeq2 normalized counts. SE is the standard error.

### miRNA target gene enrichment reveals differential regulation of regenerative and senescence-associated pathways

Using a stringent filtering strategy to identify targets of miRNAs downregulated in DPN (see Methods), we identified 1,454 target genes. Enrichment analysis of downregulated miRNA target genes in DPN yielded several pathways from the WikiPathways, KEGG, and GO databases (**Fig. 3A, Suppl. File 3**). Using the same stringent target selection protocol (see Methods), miRNAs upregulated in DPN identified a focused set of 23 gene targets that converged with targets regulated by downregulated miRNAs (**Fig. 3B, Suppl. File 3**). Pathway enrichment analysis revealed that down- and up-regulated miRNAs were associated with largely non-overlapping biological programs, indicating coordinated but opposing regulatory roles. Most commonly, terms associated with nerve regeneration, hyperglycemia/insulin resistance/dyslipidemia, cellular senescence and apoptosis and immune cell activity, were enriched with a Benjamini-Hochberg corrected p-value < 0.05. Down-regulated miRNAs were enriched for pathways involved in nerve regeneration and plasticity, including synaptogenesis, neurogenesis, neurotrophin signaling, dendritic transport, and axon regeneration, consistent with relief of repression on pro-regenerative programs. These miRNAs also mapped to insulin and lipid metabolism, AGE–RAGE signaling, autophagy, senescence, and immune pathways, suggesting coordinated regulation of metabolic stress and neuroimmune interactions. Up-regulated miRNAs showed strong enrichment for cell cycle control, reduced proliferation, cellular senescence, and regulation of apoptosis, indicating active constraint of growth and plasticity. Additional enrichment in glucose and monosaccharide response pathways links miRNA up-regulation to metabolic signaling, with concurrent effects on immune cell differentiation. Overall, miRNA down-regulation aligns with permissive regenerative and metabolic adaptation programs, whereas miRNA up-regulation is associated with growth restraint, senescence, and immune modulation.

**Figure 3.**
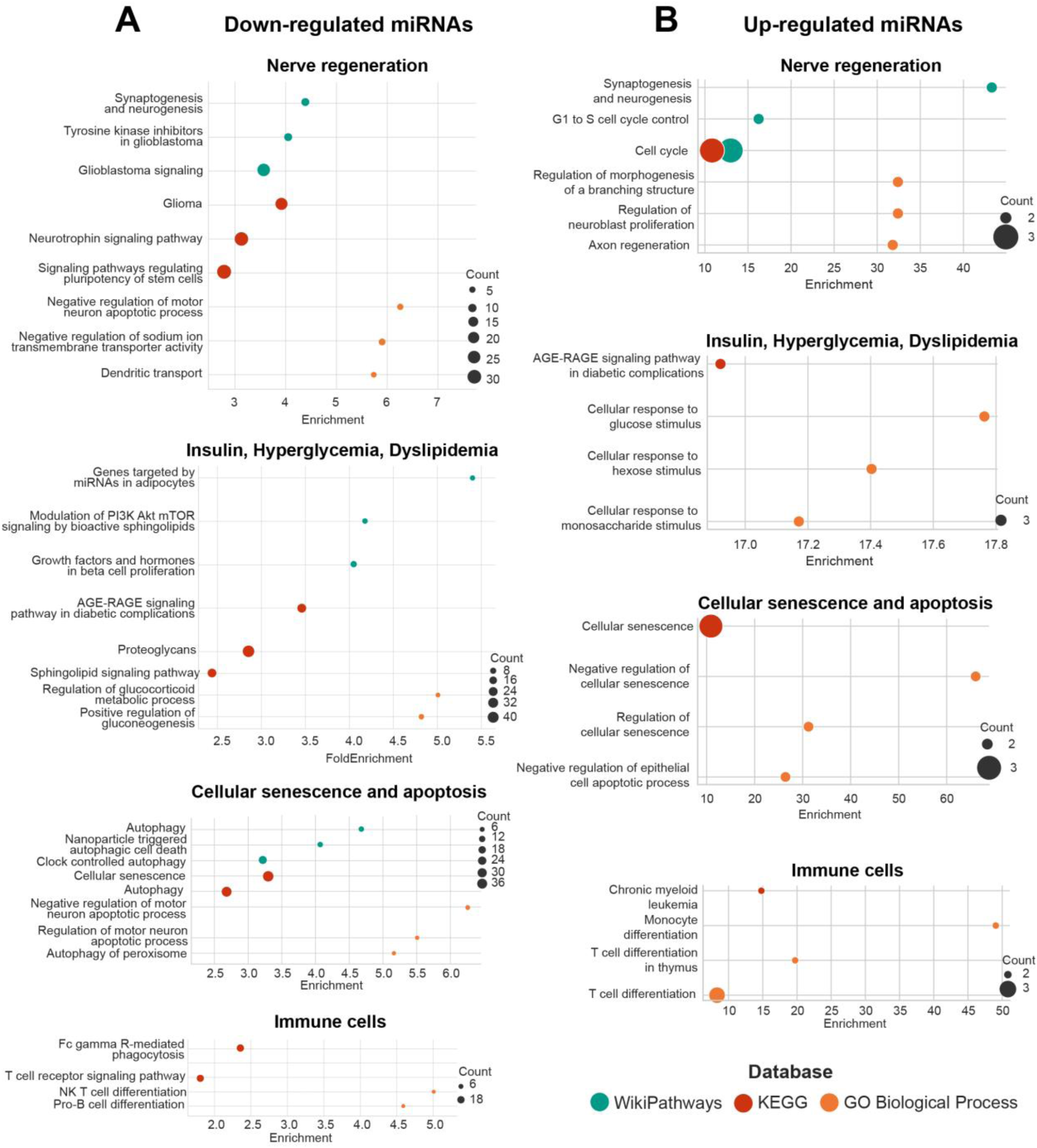
Differentially expressed miRNA Target Gene Enrichment in DPN. A) Downregulated miRNA targets top enriched terms from each database: WikiPathways, KEGG, GO Biological Process. B) Upregulated miRNA targets top enriched terms from each database: WikiPathways, KEGG, and GO Biological Process. All plotted terms have a Benjamini-Hochberg corrected p-value < 0.05.

### miR-21-5p is highly correlated with axonal loss severity and enriched in Schwann cells in DPN

Using an independent miRNA dataset comprising a small cohort of samples classified by axonal pathology, with two samples each representing normal axonal density, moderate axonal loss, and severe axonal loss as described previously ^13^, we found that several miRNAs differentially expressed between DPN and control samples were significantly correlated with axonal loss (**Fig. 4A, Suppl. File 4**). Notably, hsa-miR-21-5p (miR-21-5p) was among the most upregulated miRNAs in DPN and exhibited one of the strongest correlations with axonal loss (**Fig. 4B**), suggesting that it may be closely associated with axonal degeneration in DPN.

**Figure 4.**
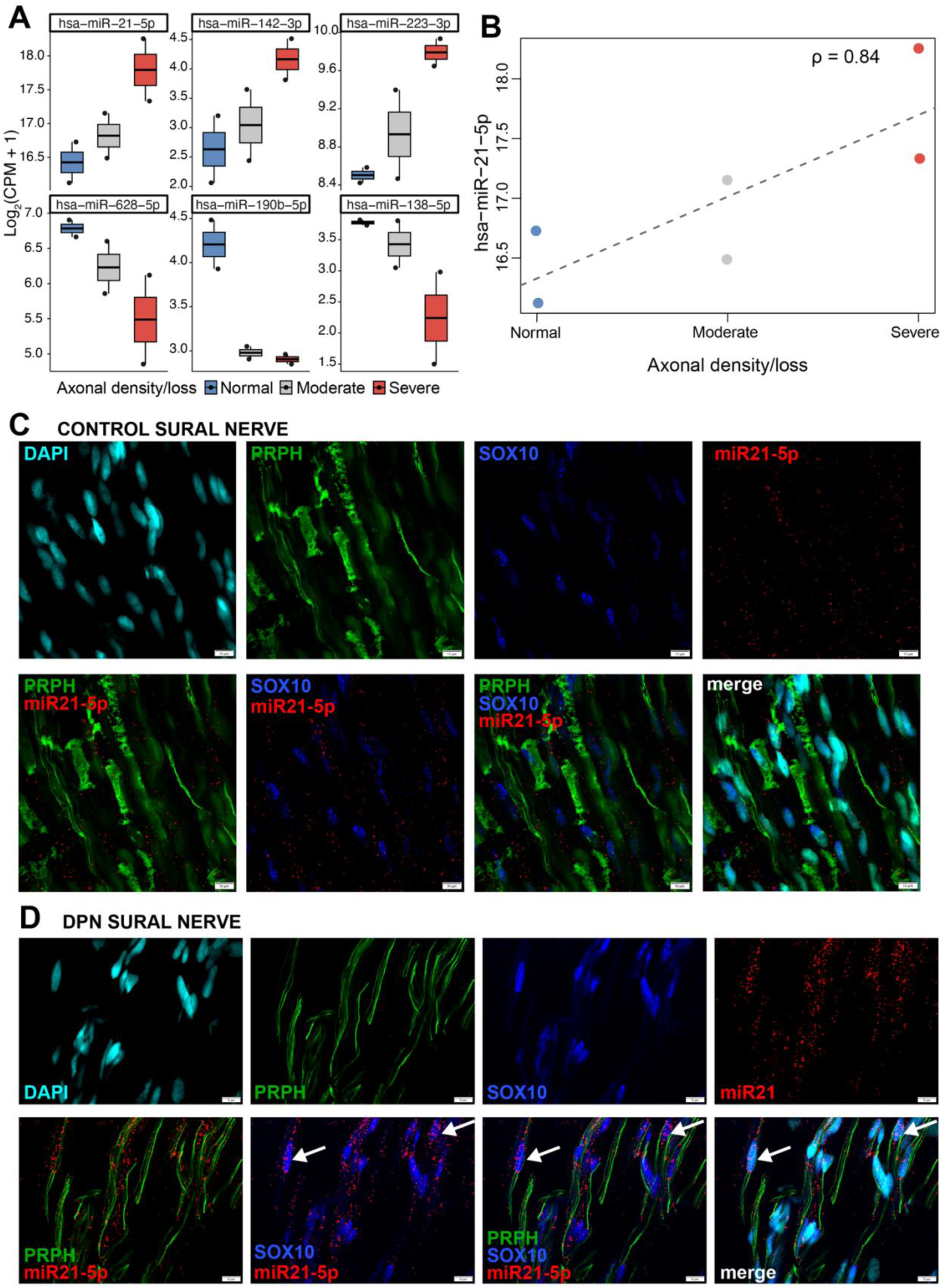
Correlation with axonal loss and spatial localization of miR-21-5p in human sural nerves. A) The six differentially expressed miRNAs between control and DPN samples showing the strongest correlations with axonal loss in an independent miRNA dataset. B) miR-21-5p exhibits a strong positive correlation with axonal loss severity. C-D) Representative fluorescence images showing miR-21-5p (red), SOX10-positive Schwann cells (blue), peripherin stained nerve fibers (PRPH, green) and DAPI (cell nuclei, cyan) in sural nerve sections from control individuals (C) and patients with DPN (D). Scale bar: 10 μm. CPM=counts per million.

Next, we sought to investigate the cellular origin of miR-21-5p dysregulation to determine which cell types contribute to its increased expression in DPN. In situ hybridization (RNAscope) revealed that miR-21-5p is broadly expressed throughout the nerve in control samples (**Fig. 4C**); however, in DPN, it exhibits increased accumulation within SOX10-positive Schwann cells (**Fig. 4D**), suggesting a potential disease-associated dysregulation in Schwann cell phenotype.

Because messenger RNA (mRNA) expression data has been previously published from the same sural nerve samples ^13^, we leveraged these matched datasets to perform integrative miRNA-mRNA analyses (**Fig. 5A**). This integrative analysis identified miR-21-5p as a highly upregulated miRNA that is negatively correlated with numerous predicted and validated mRNA targets present in the mRNA dataset, consistent with miRNA-mediated post-transcriptional repression of target mRNAs (**Fig. 5B,C, Suppl. File 5**). Target gene enrichment analysis revealed that miR-21-5p occupies a central regulatory position, targeting genes involved in axonal guidance, MAPK and Ras signaling, oxidative stress responses, and neurotrophic signaling pathways (**Fig. 5D**). Consistent with functional repression, miR-21-5p expression was inversely correlated with multiple target mRNAs, including *SOD3, PLD1, MAPK10* and *ST6GAL1* (**Fig. 5E**). These genes encode proteins that normally restrain oxidative and inflammatory stress, including extracellular antioxidant enzymes (*SOD3*), glycosylation regulators that limit immune activation (*ST6GAL1*), and lipid signaling mediators involved in inflammatory resolution (*PLD1*). Together with its enrichment in SOX10-positive Schwann cells, this suggests that elevated miR-21-5p in DPN may shift Schwann cells away from pro-regenerative and homeostatic states toward altered stress and inflammatory phenotypes, which may impair resolution of inflammation and remyelination following diabetic nerve damage.

**Figure 5.**
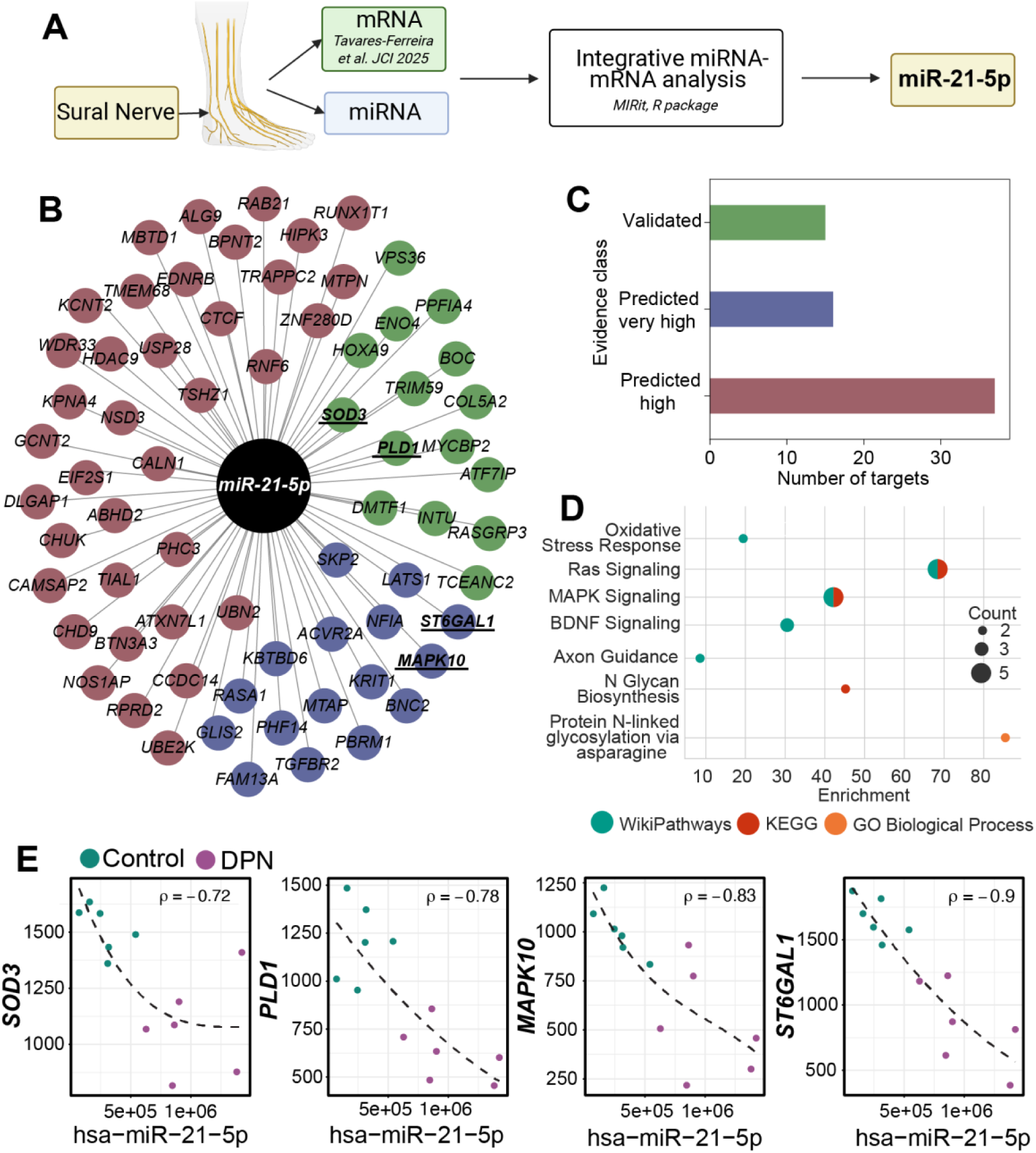
miR-21-5p target network and pathway enrichment. A) Overview of integrative miRNA-mRNA analysis focused on hsa-miR-21-5p. B) Network visualization of hsa-miR-21-5p and its predicted and experimentally supported negatively correlated mRNA targets, with node colors indicating evidence level. C) Summary of target evidence classification. D) Pathway enrichment analysis of hsa-miR-21-5p negatively correlated target genes. E) Scatter plots showing the relationship between hsa-miR-21-5p expression and representative targets *SOD3, PLD1, MAPK10 and ST6GAL1* mRNA levels in control and DPN samples. Dashed line indicates the fitted trend, with a strong inverse correlation (ρ) observed.

## Discussion

In this study, we characterize the miRNA expression landscape of human sural nerves from individuals with and without DPN. While prior studies have examined differential expression of select miRNAs in human peripheral nerves, an unbiased, high-throughput analysis of miRNA expression in human sural nerves has not been performed. Here, we provide the first comprehensive profiling of the miRNA landscape in human sural nerves in the context of DPN. We identified a total of 175 differentially expressed miRNAs, representing approximately 10% of all detected miRNAs in the sural nerve. Of these differentially expressed miRNAs, 129, or approximately 74%, were downregulated and 46, approximately 26%, were up-regulated. These miRNAs targeted genes involved in regeneration, metabolic function, senescence and apoptosis and immune cell activity. At a global level, this widespread miRNA downregulation indicates reduced post-transcriptional regulation of protein synthesis and cellular activity. This observation is consistent with prior work from our group demonstrating broad upregulation of mRNA expression in sural nerves from individuals with DPN ^13^. The extensive miRNA dysregulation observed suggests disruption of miRNA biogenesis, consistent with prior studies showing that reduced DICER and Argonaute2 (AGO2), central components of miRNA processing and effector function, leads to global decreases in miRNA abundance. Genetic deletion of Dicer led to delayed nerve regeneration in a sciatic nerve crush mouse model ^28^. In a diabetic rodent model of wound healing, Dicer expression increased by 12 days post injury in non-diabetic animals but remained unchanged in diabetic rodents ^29^. Consistent with this finding, using an in vitro wound model, increasing concentrations of Dicer small interfering RNA progressively slowed wound regeneration ^29^. Additionally, Schwann cell-specific Ago2 knockout in diabetic rodent models leads to peripheral nerve demyelination, reduced nerve conduction, and neurodegeneration ^8^. Together with the widespread miRNA downregulation observed in our dataset, these findings support a model in which diabetes may disrupt miRNA biogenesis and effector function, with downstream consequences that contribute to the pathogenesis of DPN.

To gain insight into the biological processes impacted by miRNA dysregulation, we performed target enrichment analysis of differentially expressed miRNAs. In the nerve regeneration–related category, downregulated miRNAs were enriched for neurotrophic signaling, neurogenesis, and anti-apoptotic pathways, whereas upregulated miRNAs were associated with cell cycle regulation, axon regeneration, and morphogenesis, suggesting altered regulation of neuronal survival and growth programs. Both upregulated and downregulated miRNAs converged on insulin, glucose, and lipid regulatory pathways, highlighting a close link between systemic metabolic dysfunction and peripheral nerve pathology. Previous studies have shown that RAGE activation promotes proinflammatory macrophage polarization in diabetic peripheral nerves, leading to impaired insulin sensitivity, sensory neuron atrophy, and disrupted axonal transport, while loss of RAGE shifts macrophages toward anti-inflammatory states and protects against diabetic neuropathy ^30^. Enrichment of AGE–RAGE signaling and glucose-responsive pathways supports a model in which chronic hyperglycemia and dyslipidemia reshape miRNA regulatory networks, reflecting an active but potentially maladaptive response to sustained metabolic stress. Immune-related pathways, including Fc gamma receptor–mediated phagocytosis and monocyte and T cell differentiation, were enriched in both miRNA directions, suggesting that miRNAs fine-tune immune activation states rather than simply promoting or suppressing inflammation. Interestingly, a previous study showed that chronic hyperglycemia in type 2 diabetes impairs monocyte phagocytosis through both Fcγ and complement receptors, suggesting a defect in innate immune function that may contribute to increased infection susceptibility in diabetes ^31^. In parallel, enrichment of senescence, autophagy, and apoptosis pathways, points to miRNA regulation of cell fate decisions under chronic metabolic and inflammatory stress, consistent with emerging evidence for senescence-like states in Schwann cells that limit nerve regeneration ^32^.

In our dataset, hsa-miR-21-5p emerged as a miRNA of particular interest, as it was significantly upregulated in DPN, strongly correlated with axonal loss severity and was localized to Schwann cells in DPN. miR-21-5p has been implicated across multiple nerve injuries and pathologies ^33–40^, which demonstrate that miR-21-5p is robustly induced by nerve injury, modulated by inflammatory signaling, and functions as a context-dependent regulator of axon growth, remyelination, neuronal survival, immune activation, and neuropathic pain through pathways involving PTEN-mTOR, MAPK, and inflammatory gene networks. Functionally, miR-21-5p appears to play a context-dependent role in nerve function. Acute induction of miR-21-5p supports neurite outgrowth, axon regeneration, and Schwann cell-mediated repair through suppression of inhibitory targets and activation of PTEN-mTOR signaling ^36^. However, in chronic injury or disease states, sustained miR-21-5p expression shifts toward maladaptive outcomes, including persistent inflammation, fibrotic signaling, and impaired regeneration ^41^. In our DPN nerve samples, prolonged miR-21-5p upregulation in Schwann cells may therefore amplify inflammatory and oxidative stress pathways, contributing to Schwann cell dysfunction and failed nerve repair, consistent with its strong correlation with axonal loss severity.

A recent study characterized circulating miRNA expression in type 2 diabetes and DPN and found that miR-375 levels were similar between diabetes and healthy controls but were significantly reduced in patients with DPN, supporting a specific association with neuropathy rather than diabetes itself ^42^. In our sural nerve dataset, miR-375-3p was unchanged and miR-375-5p was not detected, suggesting tissue specific regulation and that circulating miR-375 alterations may reflect systemic or non-neuronal sources rather than intrinsic changes within peripheral nerve tissue.

Overall, our study demonstrates that miRNAs are significantly dysregulated in the sural nerves of patients with advanced DPN and identifies robust associations with pathways related to oxidative stress, metabolic dysfunction, immune activity and nerve regeneration. While the present work focuses on defining these disease-associated miRNA signatures, it also establishes a strong foundation for future mechanistic and translational studies. Integrating functional validation and high-throughput spatial or single-cell miRNA approaches will enable precise mapping of these regulatory programs to specific cell types and microenvironments within the nerve. Together, these findings underscore the central regulatory role of miRNAs in DPN pathophysiology and highlight their promise as actionable biomarkers and therapeutic targets.

## Author contributions

DT-F designed the study. DT-F performed miRNA extractions for small RNA sequencing. SAG developed analysis pipeline and analyzed miRNA sequencing data including target enrichment analysis. RJ and VP performed RNAscope and IHC experiments. ECM performed integrative miRNA-mRNA analysis. SMR collected control sural nerves. DT-F and SAG wrote the manuscript. All authors read and edited the paper.

## Acknowledgments

We thank the patients for participating in this study. Funding: National Institutes of Health grant U19NS130608 (DT-F)

## Supplementary Files

**Suppl. Table 1:** Additional metadata

**Suppl. File 1**: Filtering Terms for Gene Enrichment Terms

**Suppl. File 2**: miRNA counts and differentially expressed miRNAs

**Suppl. File 3**: miRNA Target Gene Enrichment Terms Sorted

**Suppl. File 4**: miRNA and axonal loss severity spearman correlation

**Suppl. File 5**: miR21-5p negatively correlated targets from bulk mRNA-seq and target gene enrichment analysis

