## Supplementary material for "Small RNA sequencing of human sural nerves identifies widespread microRNA dysregulation and Schwann cell-localized miR-21-5p in diabetic peripheral neuropathy": Suppl. File 1

**Filtering Terms for Gene Enrichment Terms**

**Inclusionary Criteria**

Interleukins: IL,Interleukin,interleukin

MAPK: MAPK,MAP kinase,MAP Kinase,MAP,mitogenic

TGF-β/SMAD: TGF,transforming growth factor,Transforming Growth Factor,Transforming growth factor,SMAD,Smad

Immune Cell Activity: T cell,T Cell,phag,B cell,B Cell,PML,leuk,Leuk,monocyte,Monocyte,lyso,Lyso,taxis

AGE/RAGE: AGE,Advanced glycation,Advanced Glycation,advanced glycation

TNF-α: TNF,tumor necrosis factor,Tumor Necrosis Factor,Tumor necrosis factor

NTRK: NTRK,Neurotrophic,neurotrophic

Nerve Related Regeneration: sens,Sens,cytoskeleton,Cytoskeleton,Cell cycle,cell cycle,microtubule,Microtubule,dendritic spine,Dendritic spine,NGF,nerve growth factor,Nerve growth factor,Nerve Growth Factor,filopodium,Filopodium,podium,ruffle,synap,growth,Growth,Synap,Neuro,neuro,growth cone,Growth cone,Growth Cone,sodium,Sodium,potassium,Potassium,axon,astro,Astro,dendri,Dendri,vesicle,Vesicle,stem cell,Stem cell,Stem Cell,Schwann,migration,Migration,glia,Glia,glio,Glio,organelle,Organelle,Actin,actin,organization,Organization,project,Project,motil,Motil,cytosis,transport,Transport,sprout,Sprout,filament,Filament,Calcium,calcium,action potential,Action potential,Action Potential, ion,Ion,long-term,Long-term,Long-Term

Cell Death: apopto,Apopto,phag,senescence,Senescence

Extracellular Matrix/Fibrosis: adhe,Adhe,fiber,endothelial,cell-cell,Cell-cell,fibro,Fibro,matrix,Matrix,ECM,mesenchym,Mesenchym,connective,extracellular,Extracellular,junction,Junction,FGF

Insulin/Hyperglycemia/Dyslipidemia: beta cell,Beta cell,diabet,Diabet,Oxidative,oxidative,IRS,IGF,insulin,Insulin,hypoxia,fat,Fat,adip,Adip,gluc,Gluc,oxygen,Oxygen,lipid,Lipid,Chemical stress,chemical stress,nutrient,Nutrient,starvation,Starvation,starve,Starve,carbohydrate,Carbohydrate,glyc,Glyc,sacc,ose

**Exclusionary Criteria**

Nerve Related Regeneration: mesenchy,Mesenchy,organ formation,gonad,Gonad,cancer,aort,Aort,Cancer,AGE,sex differentiation,gastr,Gastr,ossif,Ossif,leukocyte,Leukocyte,MAPK,skelet,Skelet,muscle,Muscle,artery,Artery,hemat,Hemat,nephr,Nephr,renal,Renal,kidney,Kidney,derm,Derm,hair,Hair,eye,Eye,heart,Heart,ventri,Ventri,atrio,Atrio,atria,Atria,outflow,Outflow,epithel,Epithel,uter,Uter,ureter,Ureter,gland,Gland,cardi,Cardi,transforming growth factor,appendage,limb,lung,Lung,Transforming growth factor,Transforming Growth Factor,fibroblast,Fibroblast,face,Face,trachea,Trachea,virus,Virus,cartilage,Cartilage,bone,Bone,Fcgamma,hepat,Hepat,beta cell,Beta cell,starvation,Starvation,osteo,Osteo,insulin,Insulin,phag,Phag,trabecula,Trabecula,luteini,Luteini,ovul,Ovul,Ovar,ovar,Interleukin,interleukin,leukemia,endothelial cell chemotaxis,adhe,Adhe,fiber,endothelial,fibro,Fibro,matrix,Matrix,ECM,mesenchym,Mesenchym,connective,extracellular,Extracellular,junction,Junction,FGF,T cell,T Cell,phag,B cell,B Cell,PML,leuk,Leuk,monocyte,Monocyte,lyso,Lyso,chond,Chond,metria,prolactin,Prolactin,parasite,Parasite,oocyt,Oocyt,renin,Renin,aldosterone,Aldosterone,reabsorption,Reabsorption

Immune Cells: Interleukin,interleukin,leukemia,endothelial cell chemotaxis
