## Supplementary material for "Small RNA sequencing of human sural nerves identifies widespread microRNA dysregulation and Schwann cell-localized miR-21-5p in diabetic peripheral neuropathy": Suppl. Table 1

**Table S1. Additional metadata**

| Patient | Age | Sex | Diabetes | DPN | Experiment | Axonal density/loss severity |
|---|---|---|---|---|---|---|
| S10 | 62 | Female | Type 2 | Yes | Correlation with axonal severity | Moderate |
| S20 | 57 | Male | Type 2 | Yes | Correlation with axonal severity | Moderate |
| S21 | 39 | Male | Type 2 | Yes | Correlation with axonal severity | Severe |
| S22 | 63 | Female | Type 2 | Yes | Correlation with axonal severity | Severe |
| S23 | 51 | Male | Type 2 | Yes | Correlation with axonal severity | Normal |
| S24 | 58 | Male | No | No | Correlation with axonal severity | Normal |
| DPN1 | 31 | Male | Type 2 | Yes | miRNAscope | - |
| DPN2 | 79 | Male | Type 2 | Yes | miRNAscope | - |
| C1 | 36 | Male | No | No | miRNAscope | - |
| C6 | 60 | Male | No | No | miRNAscope | - |
